# Osmotic adaptation rather than stress response: A time-resolved proteomic analysis of PEG-induced water limitation in *Phytophthora cinnamomi*

**DOI:** 10.64898/2026.08.30.747438

**Authors:** Leann Seesom Vinson, Trevor Loo, Samarth Kulshreshtha, Renwick Charles Joseph Dobson, Claudia-Nicole Meisrimler

## Abstract

Water availability is critical for plants and their microbial communities, including pathogens. The plant pathogen *Phytophthora cinnamomi* persists in soils with fluctuating moisture, yet cellular responses to water limitation remain poorly understood in *Phytophthora* and oomycetes more broadly. Although we recently characterized the proteomic response of *P. cinnamomi* to NaCl-induced osmotic and ionic stress, its response to PEG-mediated water limitation remains poorly understood, leaving a critical gap in our understanding of drought-relevant stress adaptation. Here, we quantified mycelial growth and profiled time-resolved proteome dynamics of *P. cinnamomi* during polyethylene glycol (PEG-3350)-treatment, simulating moderate water limiting conditions. Treatment with 5% PEG-3350 enhanced radial mycelial growth relative to controls, with no early growth inhibition observed. Label-free proteomics identified 1,097 protein groups, with 880 proteins shared between conditions and an asymmetric abundance profile dominated by decreasing protein abundance over time. Only a small subset of proteins increased, mainly enzymes involved in redox buffering (e.g., thioredoxin and glutaredoxin-like proteins) and mitochondrial/metabolic regulation (e.g., alternative oxidase) and mitochondrial/metabolic regulation. Hierarchical clustering revealed a potential three-phase temporal program: early translational and regulatory remodeling (1–6 HPT), sustained metabolic adjustment (6–12 HPT), and delayed engagement of redox and proteostasis functions (12–24 HPT). Network analysis demonstrated that redox-associated function was integrated throughout this adaptation, with individual clusters further specialized by cofactor preference (NADP-versus NAD-dependent enzymes) and distinct metabolic roles (malate dehydrogenase, CoA-ligase activity). This coordinated, multi-phase reorganization sustained mycelial growth despite moderate osmotic stress, indicating that *P. cinnamomi* employs active proteomic adaptation rather than passive stress tolerance. These findings reveal the cellular mechanisms underlying drought persistence in this invasive pathogen and suggest molecular targets for disease management under water-limited conditions.

## Introduction

*Phytophthora cinnamomi* (*P. cinnamomi*) is an invasive soilborne oomycete with an exceptionally broad host range and major impacts on agriculture, horticulture, and native ecosystems [1,2,2–4]. Its broad distribution and persistence across diverse climates suggest that *P. cinnamomi* tolerates periods unfavorable for infection and growth, including fluctuating soil moisture [1,5]. Disease outbreaks often follow environmental shifts that enhance pathogen virulence, survival, or dispersal [6–9]. Understanding how *P. cinnamomi* persists during environmental stress and resumes proliferation under favorable conditions is therefore central to explaining its ecological success and epidemiology.

Water availability is a defining constraint in soil environments. Under current climatic patterns, with more frequent drought episodes and intermittent re-wetting it shows increasing variability. For soilborne pathogens, reduced water availability can limit turgor maintenance, alter metabolism, constrain macromolecular processes, and change the balance between growth and homeostasis [9–11]. Current understanding of drought-disease interactions remains largely host-centered. In contrast, the cellular and molecular mechanisms by which *Phytophthora* species, particularly *P. cinnamomi*, respond directly to water limitation in the absence of a host remain comparatively underexplored[12–14]. This knowledge gap is critical because understanding pathogen stress responses independent of host interactions reveals the physiological constraints and adaptive capacity that enable survival in fluctuating soil environments. Specifically, it is necessary to determine whether drought-like conditions impair *P. cinnamomi* physiology, triggering passive stress tolerance, or instead activate regulated survival programs that support persistence until conditions improve. Distinguishing between these mechanisms is essential for predicting pathogen behavior under climate-change-driven moisture variability and for identifying vulnerabilities exploitable by disease management strategies.

Polyethylene glycol (PEG) is a standard chemical treatment to mimic osmotic stress for studying water-limitation responses. It is a non-penetrating osmoticum that reduces water availability without introducing ionic toxicity [15,16,16]. This experimental approach is crucial for distinguishing the effects of osmotic stress from those of ionic stress, which we recently investigated using NaCl treatment [17].While Natural drought involves water limitation alone salt stress (e.g., NaCl exposure) combines osmotic and ionic components. By using PEG, we can distinguish the proteomic adaptations specific to water scarcity from those triggered by ion toxicity distinction rarely examined in oomycetes. PEG-induced stress in cell wall-containing eukaryotes commonly induces osmotic adjustment, metabolic reprogramming, ROS detoxification, and growth reduction. In filamentous fungi, conserved HOG/MAPK and cell wall integrity pathways coordinate glycerol accumulation, stress signaling, and maintenance of cellular integrity [18,19]. Understanding these conserved pathways in *P. cinnamomi* under osmotic stress provides a mechanistic foundation for predicting how this pathogen responds to drought independent of ion-mediated stress.

Here we address the knowledge gap of how *P. cinnamomi* responds to water limitation at the protein level, independent of the host. Growth assays were accomplished over 0–96 h post PEG treatment (HPT). Furthermore, time-resolved proteomics for timepoints 1,6,12 and 24 HPT, the temporal protein showed significant changes in protein abundance. Multivariate analysis identified distinct response patterns between treatments, while differential abundance testing pinpoints key stress-responsive proteins. Finally, network-based functional interpretation revealed underlying biological pathways and protein interactions. The present proteomic analyses of osmotic stress in *P. cinnamomi* are a first step for identifying stress-adaption with applied disease-management potential and for defining the molecular mechanisms that enable these highly invasive phytopathogens to persist under fluctuating water availability.

## Materials and methods

### *P. cinnamomi* cultivation and growth conditions

*P. cinnamomi* strain NZFS3750 (SCION culture collection) were cultivated on 10% clarified V8 (cV8) agar [20]. Prior to experimentation, mycelial plugs were taken from the leading edge of hyphal growth and subcultured onto 10% cV8 broth and incubated in the dark at room temperature for 4 days. The resulting mycelium was then collected or maintained in liquid media.

### Experimental design and sample preparation

For the mycelial growth assay, five technical replicates were used for each time point for 2.5% PEG-3350, 5% PEG-3350 and control samples, resulting in a total of 25 samples per group (a total of 75 samples). For protein quantification, experiments comprised of three biological replicates and two technical replicates per sample at each time point (six samples in total). Time points were set at 0, 1, 6, 12, and 24 hours post-treatment (HPT) for both control and 5% PEG-treated groups, resulting in a total of 30 control samples and 24 treated samples. Controls were untreated cultures collected at the same time points to ensure differences could be attributed to PEG stress. Colony radial expansion was quantified using image analysis (ImageJ, NIH). Photographic images of plates were acquired at each timepoint with a 10 mm scale reference included in each image to calibrate measurements. Colony boundaries were traced manually, and total colony area (mm²) was calculated relative to the scale. Two-way ANOVA was performed in Perseus (version 2.0.10.0) with osmotic treatment and time as fixed factors, followed by post-hoc pairwise comparisons (Bonferroni correction, α = 0.05) to determine significant differences in growth between control and PEG-treated samples at each timepoint. Randomization of samples was not relevant, as cultures were grown and harvested in parallel under identical conditions, and treatment assignment was based on the addition of PEG. For samples undergoing proteomics, the replication strategy involving both biological and technical replicates was chosen to ensure robustness at both the sample and instrument levels. This sample size and replicate design are consistent with proteomic standards for oomycete stress studies. It ensures sufficient statistical power to detect moderate effect sizes to illustrate significant differences. Mass spectrometry data were processed in Perseus (version 2.0.10.0) [21]. Proteins with ≥80% valid values per group were retained, missing values were imputed from a normal distribution (width = 0.3, downshift = 1.8), and LFQ intensities were log_−_ transformed. Principal Component Analysis (PCA) and Uniform Manifold Approximation and Projection (UMAP) were performed in Perseus and MATLAB, respectively), to assess sample clustering, detect outliers, and evaluate the consistency of biological replicates across treatments. Differential abundance analysis was conducted using ANOVA and two-sample t-tests with false discovery rate (FDR) correction (≤0.05). A multiple-sample test (one-way ANOVA, FDR ≤0.05) was first applied to identify proteins showing significant variation across time points and treatments. Subsequently, volcano plots were generated from two sample t-tests with permutation-based FDR correction (≤0.05) to visualize the pairwise differences between the control and the treated samples at each time point. Proteins were further filtered and subjected to z-score normalization to standardize the protein intensities for clustering and heatmap visualization [21]. Each treated time point was then normalized to its corresponding control, and hierarchical clustering was performed using Pearson’s correlation as the distance metric. Heatmaps were generated to visualize the protein expression patterns.

### Protein extraction and quantification

*P. cinnamomi* mycelium was weighed and ground into a fine powder in liquid nitrogen. The powder was resuspended in extraction buffer (50 mM Tris-HCl, pH 7.6, 150 mM NaCl, 2% SDS, 8 M urea, Milli-Q water) supplemented with a protease inhibitor cocktail (cOmplete™, MilliporeSigma), incubated on ice for 1 hour, and vortexed periodically. After incubation, samples were centrifuged at 13,000 rpm for 15 min at 4 °C, and the supernatant was collected. For protein quantification, the supernatant was diluted 1:4 (sample: Milli-Q) and dispensed into a 96-well plate containing Pierce™ 660 nm Protein Assay Reagent (Thermo Fisher) and Ionic Detergent Compatibility Reagent (Thermo Fisher), alongside a BSA standard curve. Protein concentrations were determined as per manufacturer’s instructions. Absorbance at 660 nm was measured, and concentrations calculated in GraphPad Prism 9 (v9.5.1).

### Polyacrylamide gel electrophoresis (PAGE), staining and destaining

Mycelial protein samples were mixed with 4x LDS sample buffer (Thermo Fisher Scientific) containing 10 mM DTT and heated at 90 °C for 10 min, followed by centrifugation at 15,000 G for 10 min. A total of 150 µg of *P. cinnamomi* mycelial protein was loaded into each well of a Bolt™ 4–12% Bis-Tris gel (1.0 mm, Mini Protein Gels, Thermo Fisher Scientific). Electrophoresis was accomplished in 1x MES buffer for 30 s to 2 min. at 200 V, until the protein entered the resolving gel. Following electrophoresis, gels were rinsed with Milli-Q water and stained with Coomassie Brilliant Blue solution (Brilliant Blue, (NH) SO, Milli-Q water, 85% phosphoric acid) for 5 min at room temperature. Gels were agitated during staining and subsequent washes until background destaining was sufficient. The completed stained protein bands were excised with a scalpel under Milli-Q water and transferred to tubes containing 80% acetonitrile. Tubes were vortexed for 15 min, and the destaining step was repeated until the gel pieces appeared clear. Gel pieces were then incubated in fresh destaining solution for 5 min at room temperature, after which the acetonitrile was removed, and the tubes were dried in a SpeedVac for 4–7 min.

### Reduction, alkylation, trypsin digestion and protein extraction*-*

Reduction and alkylation were performed as described in [22] with minor modifications. Gel pieces were incubated in reducing solution (1× ammonium bicarbonate [ABC], 10 mM DTT) at 45 °C for 1 hour with periodic vortexing. The solution was replaced with 80% (v/v) acetonitrile, vortexed until the gel pieces turned white, and the supernatant was discarded. Gel pieces were dried in a SpeedVac for 4-5 min. For alkylation, gel pieces were incubated in alkylation solution (1× ABC, 20 mM iodoacetamide) at room temperature in the dark for 30 min. The solution was removed and replaced with 80% (v/v) acetonitrile, and the dehydration step was repeated twice. Finally, the acetonitrile was removed, and gel pieces were dried in a SpeedVac. Proteolytic digestion was initiated by adding 50 µL of digestion solution containing Trypsin-Ultra™ (New England Biolabs, P8101S) in Trypsin-Ultra reaction buffer (New England Biolabs, B8101S), diluted with 1× ABC to achieve a final trypsin concentration of 2% (w/w relative to protein). Gels were fully covered with the digestion solution for ∼30 min to allow enzyme penetration; excess solution was removed and replaced with 1× ABC. Digestion was carried out overnight (16 h) at 37 °C. Peptides were recovered by sequential extraction. First, gels were incubated with 5% formic acid in 40% (v/v) acetonitrile, sonicated for 2 min, and the supernatant collected. A second extraction was performed with 0.1% formic acid in 80% (v/v) acetonitrile, with sonication for 2 min, until the gel pieces turned white. All supernatants were pooled and lyophilized for 24 h prior to LC-MS/MS analysis.

### Liquid chromatography-mass spectrometry^2^ (LC/MS-MS)

Lyophilized samples were briefly spun, reconstituted in 40 µL 0.1% formic acid in 3% acetonitrile, vortexed, and sonicated for 5 min to enhance solubility. After centrifugation (15 min, 4 °C), supernatants were transferred to HPLC vials for analysis. A maximum injection of 4 µL of sample were loaded onto the enrichment trap and eluted into the 75 μm x 50 cm analytical column in a linear gradient from 3–30% acetonitrile over 60 minutes. The digested peptides were separated by their affinity for water on a reverse-phase C18 column and analyzed by mass spec using a Top10 method where a survey (MS1) scan was performed and the top 10 most intense peptide ions within a specified range were fragmented to obtain their sequence information for identification (MS2). The process was repeated for every scan cycle in the chromatographic run and additional settings enable the mass spec to ignore repeating ions within a specified time window. This allowed lower abundance ions to be targeted and improve overall coverage of the sample. In all experiments, MS1 scans were acquired over a mass range of 370–1,700 m/z with detection in the Orbitrap mass analyzer at a resolution setting of 70,000. Fragment ion spectra produced *via* higher-energy collision-induced dissociation (HCD) collision cells were acquired with a resolution setting of 17,500. Dynamic exclusion conditions were optimized according to observed chromatographic peak width (typically 6s). Details for chromatography and MS settings are listed in Supplementary table 1.

### Data processing and statistical analysis

MS data were processed using MaxQuant (v 2.4.13.0) using Andromida search engine for label-free quantification (LFQ). Peak lists were generated directly from raw Orbitrap files with default settings (de-isotoping and charge state assignment included no additional smoothing applied). Searches were performed against the combined UniProtKB and NCBI *Phytophthora* species FASTA databases (February 2024 release) supplemented with the MaxQuant common contaminants database. Protein identification was accepted at a false discovery rate (FDR) ≤ 0.05 at the protein level, as determined by target-decoy database searching, ensuring high-confidence identification of protein groups. Considerable peptide modifications included oxidation (M), acetylation (N-term), carbamidomethylation (C), and phosphorylation (STY), with Trypsin/P used for digestion, with max. missed cleavages set to 2. Label free quantification was selected, and all settings default. The precursor mass tolerance was set to 20 ppm (first search) and 4.5 ppm (main search), and fragment mass tolerance to 20 ppm. Other settings not mentioned were kept at default.

### Gene Ontology (GO) Annotation and Categorization

GO analysis was performed using protein lists derived from both differential abundance analysis (*via* volcano plots) and hierarchical clustering. Due to the limited availability of categorized GO terms for many *P. cinnamomi* proteins in the public databases, annotation was conducted manually. Protein abundance matrices from Perseus, containing significantly regulated proteins at 1, 6, 12, and 24 HPT, were exported and protein identifiers (GenBank accession numbers) were converted to UniProtKB identifiers using the UniProt ID mapping tool (https://www.uniprot.org/id-mapping). For proteins that did not yield a UniProtKB match, FASTA sequences were extracted and queried against the UniProt database using BLASTP to identify the homologous protein identity. GO terms (Biological Process [BP], Molecular Function [MF], and Cellular Component [CC]) were then manually assigned based on the closest functional homologs. Finally, GO annotations were grouped and categorized in GraphPad Prism 10.5.0 according to their respective ontology domains to visualize the distribution of functions across differentially abundant proteins.

### Network analysis

Network analysis was restricted to proteins showing significant abundance changes across timepoints and treatments. Proteins passing the multiple-sample ANOVA significance threshold (FDR ≤ 0.05) were selected for functional annotation and network analysis. Since *P. cinnamomi* is not represented in the STRING database (v12.0), interaction analysis was conducted using homologous proteins from *Phytophthora infestans* as reference IDs. Protein sequences corresponding to PEG-responsive *P. cinnamomi* proteins were queried against the *P. infestans* proteome using BLASTp. For each query, the highest-scoring BLAST hit (based on bit score and E-value) was selected as the corresponding *P. infestans* homolog. The matched *P. infestans* protein identifiers were then retrieved directly from the STRING database and used for downstream interaction and enrichment analyses. Validated protein identifiers were imported into Cytoscape (3.10.4) using the STRING app to generate a protein-protein interaction network. Interactions with a combined confidence score ≥ 0.7 were retained. Functional enrichment for Gene Ontology (GO) terms, including molecular-function categories associated with redox metabolism and oxidative stress, was performed within the STRING environment and visualized in Cytoscape. Node attributes were used to encode cluster identity (color), degree or centrality metrics (size), and inter-cluster bridging roles (border thickness). The network was visualized using the yFiles Organic layout, enabling the separation of functional modules and clear identification of proteins directly and indirectly associated with stress regulation.

## Results and Discussion

### Physiological response to Osmotic Stress

To assess physiological responses to osmotic stress, mycelial radial growth was quantified on 10% V8 agar over 0–96 h post-treatment (HPT). Two-way ANOVA identified a significant treatment by time interaction (p < 0.0001), indicating that the effect of PEG on radial growth differed across the experimental period. Early timepoints (24–48 HPT) showed no significant effect of PEG concentration on growth rates, but by 72 HPT, both 2.5% and 5% PEG treatments resulted in significantly enhanced mycelial expansion relative to untreated controls (p < 0.0001; Fig 1A–D). Post-hoc comparisons revealed that at 72 HPT, 5% PEG-treated colonies were 429 units larger (approx. 30% increase in radial expansion) compared to control, and this growth advantage was sustained through 96 HPT where treated colonies measured 507 units larger than controls (approx. 23% increase). Colony morphology remained normal under both PEG concentrations, with no visible hyphal deformation or discoloration observed. This sustained and significantly enhanced growth under PEG exposure is notable, as it contrasts with reports in other *Phytophthora* species where osmotic stress typically reduces mycelial expansion [23]. Notably, the combined osmotic and ionic stress imposed by 100 mM NaCl resulted in accelerated mycelial expansion rather than growth suppression, indicating that ionic toxicity does not limit *P. cinnamomi* proliferation at physiologically relevant salt concentrations [17]. The robust growth phenotype despite osmotic challenge suggests that *P. cinnamomi* does not experience acute physiological impairment from water limitations but instead employs cellular mechanisms that maintain or enhance expansion under conditions of reduced water availability.

**Fig 1:**
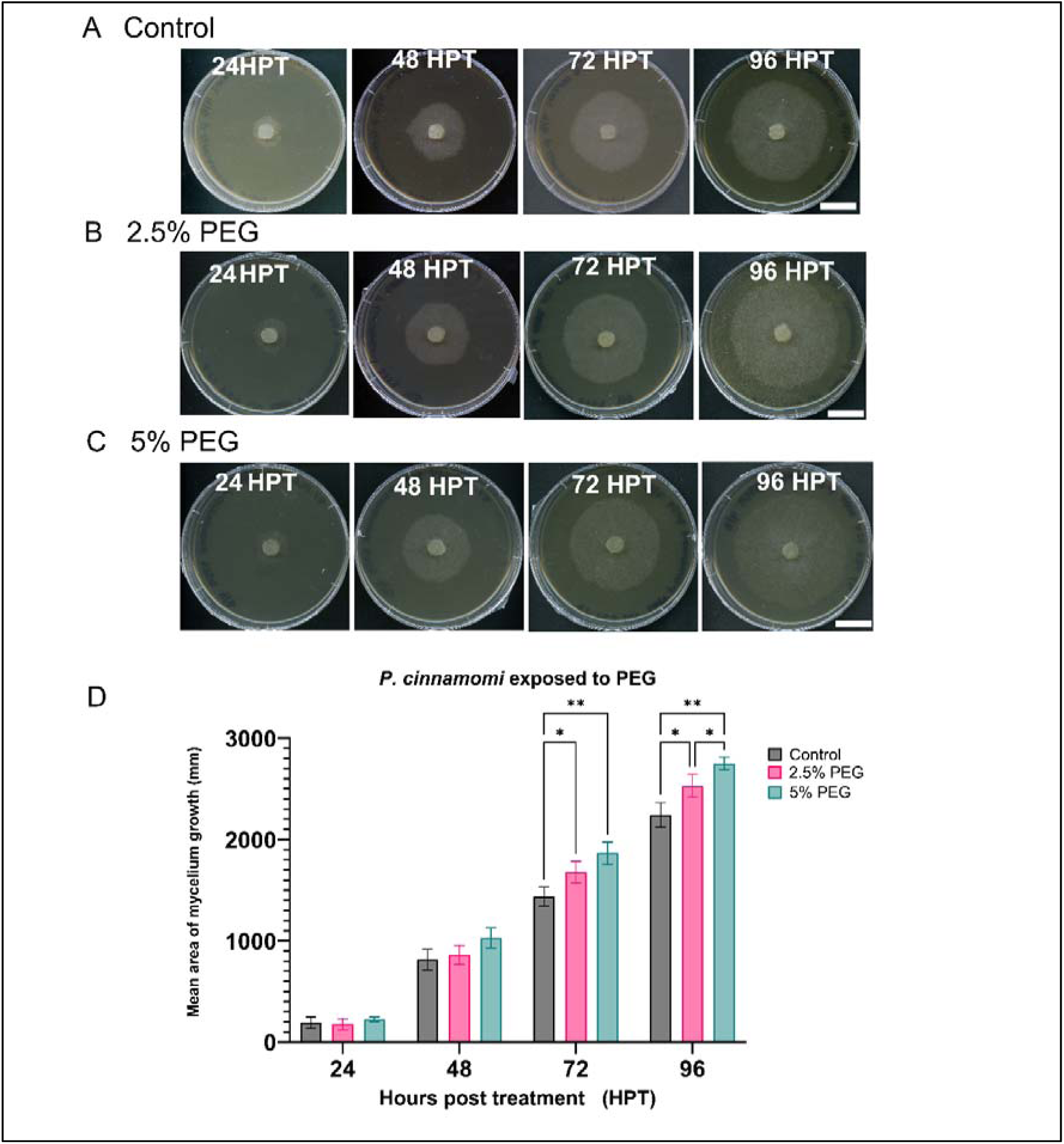
The growth rate of P. cinnamomi on 10% V8 agar plates and 10% V8 modified agar plates with 2.5% and 5% of PEG-3350 over a period of 96 hours post treatment (HPT), Scale bar: 10 mm (White block). (A) Control (cV8), (B)2.5% PEG 3350, (C)5% PEG 3350. (D) Quantification of radial mycelial growth (mean ± SD, n = 5 per time point) for control (Grey), 2.5% PEG 3350 (Pink), and 5% PEG 3350 (Blue); Statistical analysis was performed using Dunnett’s multiple comparisons test comparing each treatment to the control at each time point.Significant differences are indicated by asterisks: 48 h – “*”: p = 0.0238; 72 h – “*”: p = 0.0128, “**”: p = 0.0042;96 h – “**”: p = 0.0054. All other comparisons were not significant (p > 0.05).

### Protein coverage and data set overview

Since no physiological impairment was observed, we hypothesized that *P. cinnamomi* employs molecular mechanisms to maintain cellular homeostasis under water limiting conditions. To characterize global proteomic reorganisation under these conditions, time-resolved LC–MS/MS analysis was performed on control and 5% PEG-3350 treated *P. cinnamomi* cultures (Figure 2A). The 5% PEG concentration was selected because it showed more significant growth increase compared to the lower 2.5% concentration, ensuring detectable stress responses. However, both PEG concentrations may be considered mild osmotic stress. The analysis identified a total of 128,265 peptide-spectrum matches (PSMs), with 80,989 in the control group and 47,276 in the 5% PEG-treated group, respectively (Fig 2B, S1 Table). The total PSMs correspond to 1,977 non-redundant protein groups following database mapping from NCBI and UniProtKB. This dataset contained a comprehensive coverage of central metabolic, regulatory, and stress-associated pathways. PCA and UMAP were used to visualize overall proteomic similarity between control and 5% PEG-treated samples (Fig 2C–D) [24,25]. Both methods showed a treatment-associated shift with partial overlap between groups, indicating that PEG exposure alters global proteomic structure while a shared core proteome is maintained. Consistent separation of treatment groups across both approaches indicates that observed clustering reflects underlying biological differences rather than method-specific errors.

**Fig 2:**
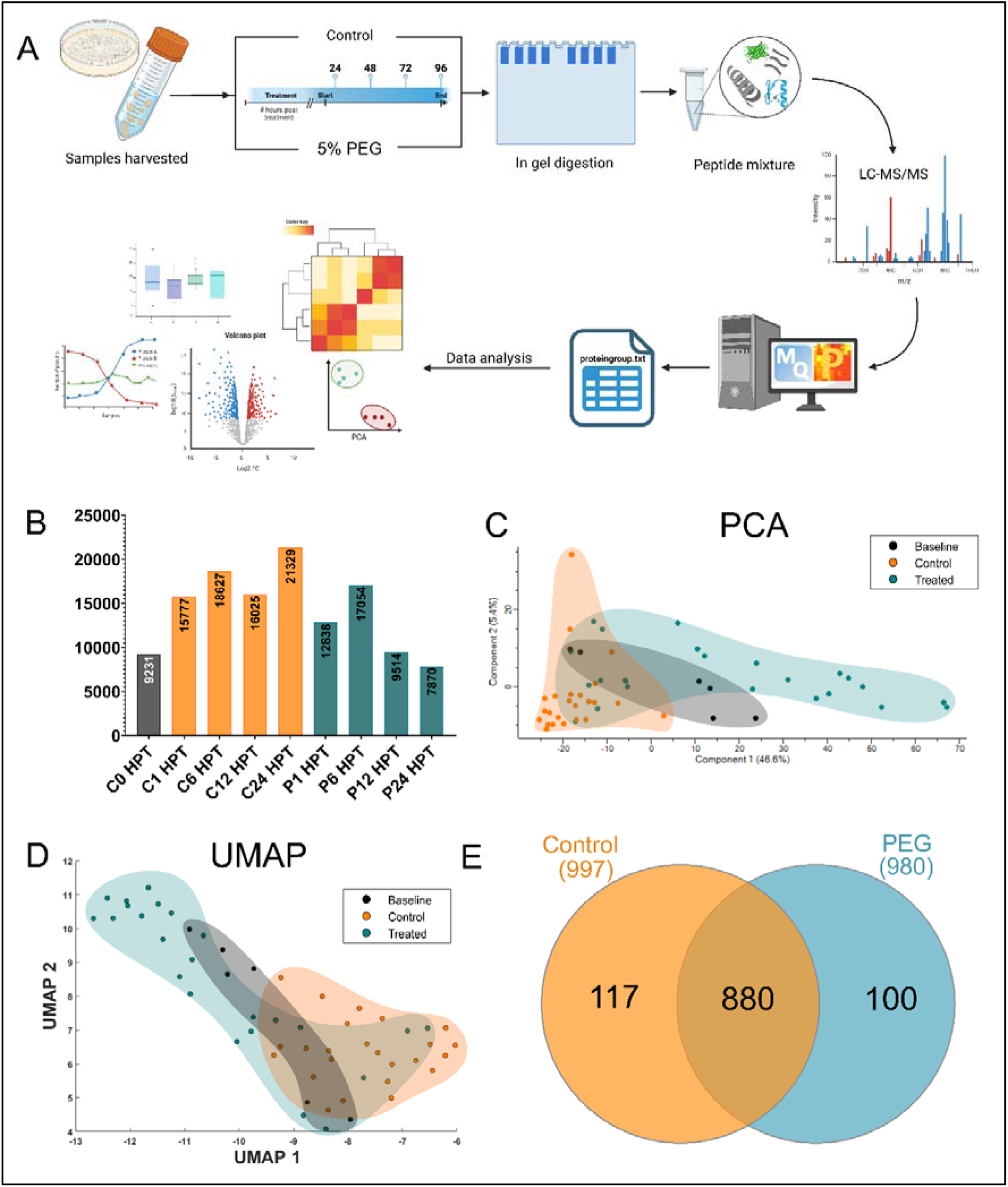
Proteomic analysis of whole mycelium proteomics exposed to osmotic stress. (A) Experimental process for *P. cinnamomi* total proteomic analysis. (B) Total number of proteins. (C) Principal component analysis (PCA). (D) Uniform Manifold Approximation and Projection (UMAP). (E) Total number of similar protein groups shared between the control and treated group, the proteins groups identified as unique to the control group (Orange) and unique to PEG treatment (Blue).

### Treatment-specific proteomes reflect growth-associated and stress-associated functional states

Comparison of protein detection across the control and PEG-treated samples revealed a large, shared core proteome, alongside a subset of proteins that were preferentially detected under one condition (Fig 2E; S1 Table). Of the total protein groups identified, 880 proteins were detected in both control and PEG-treated samples, indicating substantial overlap in core cellular functions under osmotic water limitation. In addition, 117 proteins were detected only in control samples, while 100 proteins were detected only in PEG-treated samples across the dataset.

The large proportion of shared proteins (880) suggests that PEG-induced osmotic stress does not eliminate the core metabolic capacity but instead reorganizes the detectable proteome towards stress adaptation and cellular homeostasis. The 100 proteins exclusively detected under PEG treatment represent a coordinated stress-response proteome distinct from basal cellular functions (S1 Table). In comparison, NaCl stress (100 mM) produced a similar proteome restructuring pattern with 991 proteins detected in both control and NaCl-treated samples, 51 proteins unique to control conditions, and 107 proteins exclusively detected under salt stress [17]. This proportional similarity suggests that both osmotic and ionic stresses reorganize proteomes toward specialized adaptation rather than global biosynthetic suppression.

These PEG-specific proteins associate with cellular repair and metabolic reorganization mechanisms, with prominent cysteine proteases (A0A8T1VQM1_9STRA) that potentially facilitate protein degradation and turnover essential for removing stress-damaged proteins [26,27]. DNA repair machinery, including RuvB-like helicases (KAE9246971.1) and UV excision repair proteins (KAH7488164.1) have been identified and are usually involved in protection against stress-induced DNA damage [28]. Metabolic remodeling was evident through increased representation of fatty acid metabolism enzymes (KAG6621240.1, KAG6609184.1, KAG6610728.1), alternative oxidases (KAG6610988.1) that bypasses conventional electron transport [29], and amino acid biosynthesis enzymes supporting osmolyte production for cellular osmoregulation [30].

The presence of specialized stress-responsive kinases (A0A9W7CY87, KAG6612457.1, KAG6594299.1) suggests activation of signaling cascade regulation, while molecular chaperones and protein quality control systems (A0A2P4X7A0, KAG6612208.1, KAG4056508.1) likely maintain proteostasis under the osmotic stress [31–33]. The detection of translation machinery components under PEG treatment, including eukaryotic initiation factor 4E (KAG6619349.1), translation initiation factor 3F (G5A7A3), and ribosomal proteins (KAG6602869.1, KAG6619355.1), likely facilitate stress-specific protein synthesis programs rather than general biosynthetic activity [34,35]. Notably, comparable translation machinery components, including aminoacyl-tRNA ligases and ribosomal proteins L10, L18, and S6, were similarly upregulated under 100 mM NaCl stress, suggesting that both osmotic and ionic stresses demand stress-specific protein synthesis [17]. However, the temporal dynamics diverged, while PEG-specific translation machinery appears constitutively elevated, NaCl stress triggered a coordinated temporal progression of translation capacity aligned with redox management phases. The abundance of thioredoxin-like proteins (KAG6614977.1, KAG6576379.1), glutaredoxins (KAG6617913.1), and alternative respiratory enzymes indicate enhanced antioxidant capacity and metabolic flexibility essential for surviving water limitation [36–39].

These proteins represent a proposed comprehensive stress-adaptation that enables *P. cinnamomi* to maintain cellular integrity and metabolic function during moderate osmotic challenge. While PEG and NaCl treatments both activated overlapping proteome reorganization, including translation machinery, molecular chaperones, and glutathione/thioredoxin-based antioxidant system. The mechanistic implementation differed significantly, NaCl stress uniquely induced a triphasic temporal progression (1–6 HPT, 6–12 HPT, 12–24 HPT) orchestrated through glutathione reductase as a central regulatory hub, whereas PEG-induced osmotic stress appears to engage sustained, rather than phased, antioxidant and metabolic flexibility. This suggests that while both stresses challenge cellular homeostasis through water limitation, the ionic toxicity component of salt stress demands fine-tuned temporal metabolic coordination absent under osmotic challenge alone.

### PEG exposure triggers reduction in protein abundancy

Volcano plot analysis of temporal protein abundance changes during PEG-induced osmotic stress showed pronounced asymmetry, with significantly decreased proteins substantially outnumbering significantly increased proteins across all examined timepoints (Fig 3A–D). Only 9 proteins showed significantly increased abundance, compared to 607 downregulated proteins (Fig. 3E-G). This pronounced asymmetry in the proteome response contrasts markedly with the balanced temporal proteome remodeling observed under 100 mM NaCl stress, which showed substantial protein redistribution across multiple time points (48 upregulated and 53 downregulated at 1 and 6 HPT, respectively) without the dominance of protein loss characteristic of osmotic stress [17]. This difference suggests that ionic stress promotes active proteome restructuring, whereas osmotic stress triggers more conservative downregulation. PEG-induced proteomic asymmetry has also been reported in plants; for example, wheat leaves exposed to PEG-6000 showed a predominance of decreased protein abundance, with 111 of 176 differentially expressed proteins reduced under stress [40].

**Fig 3:**
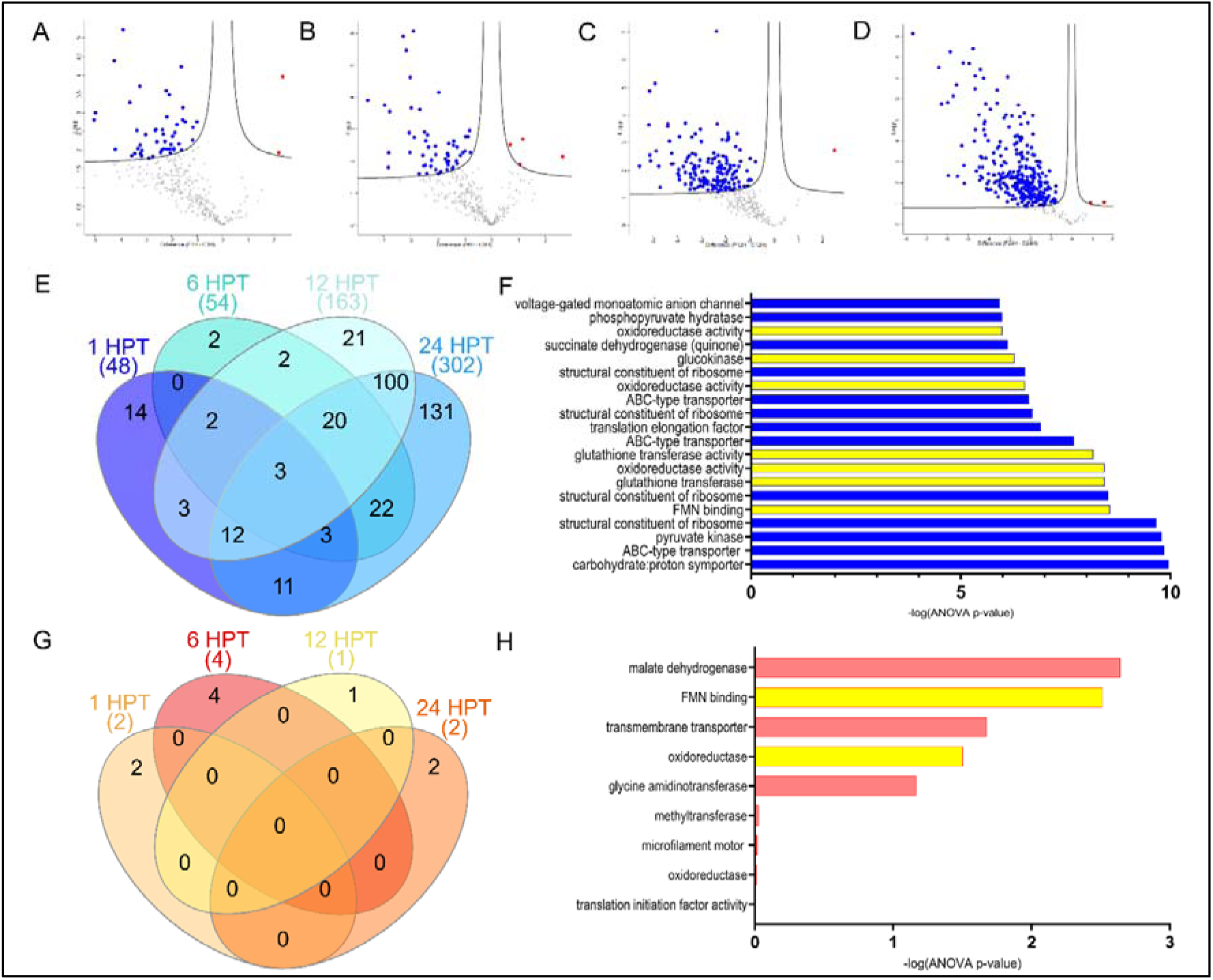
Differentially abundant proteins (DAPs) across osmotic stress time points in *P. cinnamomi*. (A-D) Volcano plots comparing control (C1, C6, C12, and C24 hours post-treatment [HPT]) and corresponding 5% PEG-3350 treated (N1, N6, N12, and N24 HPT) samples. Proteins with significantly higher or lower abundance are highlighted in red and blue, respectively. Significance was determined using Perseus default volcano plot settings (FDR = 0.05, S = 0.1). (E) Venn diagram showing overlap of low-abundance proteins across the four time points. (F) Gene Ontology (GO) enrichment analysis of the combined set of low-abundance proteins from all four time points, with enriched terms ranked by –log (ANOVA p-value). (G) Venn diagram showing overlap of high-abundance proteins among the time points. (H) GO enrichment analysis of the combined set of high-abundance proteins across all time points. For the GO analysis the word “analysis” and GO numbers were removed and are available in Supplementary table 2.

The observed decrease in protein abundance likely reflects an energy-conservation strategy during osmotic stress, as sustained synthesis and maintenance of non-essential proteins is metabolically costly under water-limiting conditions [41,42]. The reduction in protein abundance may represent a form of “proteome downsizing” where *P. cinnamomi* prioritizes the maintenance of only the most critical cellular functions while eliminating synthesis of energetically expensive but dispensable proteins. This response is consistent with the cellular economy principle, where organisms facing resource limitations shift from growth-oriented to survival-oriented metabolism [43].

During the first 24 h of PEG-induced osmotic stress, the proteome underwent a marked reorientation characterized by the coordinated downregulation of biosynthetic and translational machinery alongside selective upregulation of protective responses. Proteins with significantly decreased abundance including ribosomal proteins including 40S ribosomal protein S4 and 60S ribosomal export protein NMD3 showed decreased abundance, as did translation elongation factors including translation elongation factor 1-alpha and elongation factor 1-gamma, together with aminoacyl-tRNA ligases [44,45]. Central metabolic enzymes showing reduced abundance included glycolytic proteins: glucokinase, glyceraldehyde-3-phosphate dehydrogenase, and glucose-6-phosphate isomerase, along with multiple oxidoreductases and tricarboxylic acid cycle enzymes, collectively indicating suppression of both anabolic and catabolic pathways during the acute stress phase. (Fig. 3F, S2 Table) [46]. In contrast, the smaller set of proteins with significantly increased abundance was enriched for stress-associated functions, including FMN binding and oxidoreductase activity, consistent with selective activation of protective responses rather than broad metabolic induction (Fig. 3H, S2 Table) [47–49]. NAD-dependent redox enzymes showed prominent increases, including malic enzyme and phosphoglycerate dehydrogenase, both supporting energy metabolism and amino acid biosynthesis under osmotic stress [50]. Antioxidant defenses were bolstered by NAD(P)H:quinone oxidoreductase, type IV (A0A225WR50), which provides ROS scavenging through quinone reduction [51]. Translation machinery showed selective maintenance, with eukaryotic translation initiation factor 2 subunit 3 (A0A8T1X9Z5) increased to preserve stress-responsive protein synthesis [52]. Osmolytic capacity may be sustained through glycine amidinotransferase (A0A6A3U9T8), supporting creatine and polyamine biosynthesis for cellular protection [53].

Notably, this early proteomic reduction and selective increase of protective responses occurred despite sustained and enhanced mycelial growth on solid medium (72 HPT and 96 HPT), suggesting that *P. cinnamomi* may maintain hyphal expansion under water-limiting conditions while reducing the abundance of proteins associated with growth-related cellular processes. This apparent paradox, growth sustained despite reduced growth-associated proteins, could suggest that the proteome reorganization enables efficient resource allocation, channeling energy and biosynthetic capacity toward osmotic adaptation and survival rather than growth biomass. Interestingly, this growth-despite-downsizing phenotype under osmotic stress contrasts sharply with the salt stress response, where mycelial expansion was actively enhanced alongside coordinated upregulation of energy metabolism and biosynthetic enzymes [17]. This distinction suggests that osmotic challenge triggers defensive proteome minimization, whereas ionic stress triggers offensive metabolic activation, a fundamental difference in adaptation strategy that may reflect the distinct physiological demands of water limitation versus ionic toxicity. How mycelial expansion is metabolically sustained under these contrasting proteome states remains to be elucidated through further investigation of integrated pathway analysis and metabolic modeling.

### Growth and pathogenicity responses demonstrate temporal remodeling rather than persistent suppression

Rather than global developmental arrest, proteins associated with cytoskeletal organization and growth machinery showed selective upregulation at 6 HPT (actin filament organization, myosin complex), indicating that growth-related functions undergo dynamic remodeling rather than sustained shutdown [19,54]. Notably, this temporal reactivation of growth machinery occurred alongside the early proteomic reduction, suggesting that *P. cinnamomi* may initially suppress growth-associated protein synthesis during the acute stress phase (1–6 HPT) before selectively reengaging cytoskeletal remodeling at 6 HPT, potentially to reorient hyphal architecture under osmotic limitation. In parallel, translational machinery exhibited selective suppression: bulk translation factors (40S ribosomal proteins, elongation factors EF1-α and EF1-γ, tRNA synthetases) were significantly downregulated across all timepoints, while eukaryotic translation initiation factor 2 subunit 3 was selectively upregulated at 6 HPT, suggesting that *P. cinnamomi* employs differential translational control suppressing global protein synthesis while maintaining capacity for targeted translation of stress-response factors [45,55]. Virulence-associated proteins, including secreted RxLR effectors and metallopeptidases (Cluster H), remained at significantly suppressed but negligible abundance levels throughout the 24 h period, indicating stable deployment inhibition consistent with prioritization of immediate osmotic adaptation and cellular survival over pathogenic functions. Together, this temporal proteome architecture reveals a coordinated adaptive strategy. Growth remodeling, selective translation, and virulence suppression are sequentially orchestrated across the osmotic stress response, demonstrating that *P. cinnamomi* prioritizes immediate survival over pathogenic activity during water limitation.

### Three-phase adaptation program integrates translation, metabolism, and redox control

While differential abundance analysis captured individual protein changes, it was not resolved whether these changes follow coordinated temporal patterns. Hierarchical clustering was therefore used to group proteins with shared abundance dynamics across the PEG exposure. This analysis revealed a progressive response indicating that osmotic stress response in *P. cinnamomi* is driven by active, energy-dependent adaptation integrating translational control, metabolic restructuring, and redox homeostasis, involving selective rather than global protein synthesis changes (Fig 4A-B, S3 Table).

**Fig 4:**
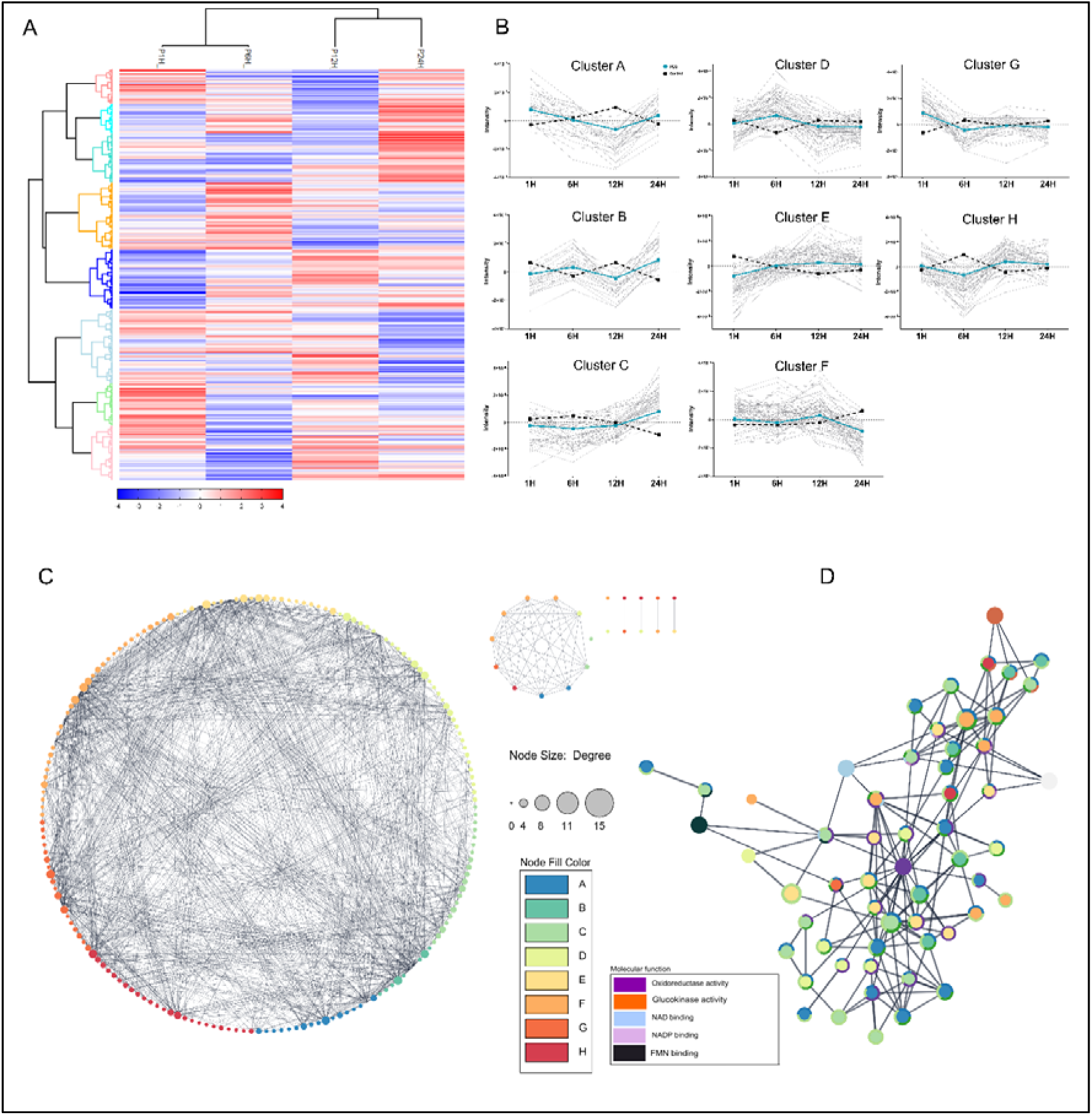
Hierarchical clustering and network analysis of *P. cinnamomi* proteins under 5% PEG-3350 exposure. (A) Heatmap of ANOVA-significant proteins (FDR ≤ 0.05), z-scored and partitioned into eight clusters (A–H) by hierarchical clustering in Perseus, shown across 1, 6, 12, and 24 h post-treatment (HPT). (B) Temporal abundance profiles for each of the eight clusters (A–H) across 1, 6, 12, and 24 HPT; grey lines represent individual proteins, with dashed black and solid teal lines indicating control and PEG-treated group averages, respectively. (C) Protein-protein interaction network of differentially abundant proteins (DAPs) mapped in STRING (combined score ≥ 0.7) and visualized in Cytoscape; node color corresponds to cluster assignment (A–H) and node size reflects degree (number of connections). (D) Subnetwork of 96 proteins associated with ROS/redox regulation, colored by cluster assignment(A: blue; B: green; C:light green; D:yellow; E:light orange; F: orange; G: Dark orange; H: red), with outline color indicating molecular function annotation (Purple: oxidoreductase activity, Orange: glucokinase activity,Blue: NAD binding,Lilac: NADP binding, Black FMN binding).

Early-phase responses (0-6 HPT) were dominated by proteins associated with RNA binding, ribosomal structure, translation initiation and elongation, and ATP-dependent regulation (S3 Table). While many proteins were downregulated globally to conserve energy, this pattern indicates active translational reprogramming with selective upregulation of stress-relevant machinery rather than complete suppression of protein synthesis, consistent with prioritization of essential stress-response transcripts [55–57]. From a systems biology perspective, this selective translational control allows the pathogen to rapidly redirect limited energy resources toward stress-adaptive proteins while minimizing metabolic burden, establishing the foundation for survival during osmotic challenges (Fig 4A-B, S3 Table). Notably, this early-phase selective translational upregulation under osmotic stress parallels the immediate translation machinery activation observed during 100 mM NaCl exposure (1–6 HPT), where eukaryotic initiation factors and ribosomal proteins were similarly upregulated [17]. However, a critical mechanistic distinction showed NaCl stress triggered direct and immediate activation of ROS-detoxifying enzymes (glutathione S-transferases, peroxidases) within this same 1–6 HPT, whereas PEG-induced osmotic stress shows minimal redox enzyme upregulation at early timepoints. This suggests that translational reprogramming is a conserved rapid response to both stress types, but the presence of antioxidant mobilization diverges based on the primary stressor (ionic toxicity versus water limitation).

Mid-stage responses (6-12 HPT) were enriched for central carbon metabolism, amino acid and nucleotide biosynthesis, and ATP-dependent transport and enzymatic activity, indicating sustained metabolic demand under continued water limitation [58–61] (S3 Table). Despite continued global protein downregulation, ribosomal and translation-associated proteins remained selectively maintained, supporting biosynthetic capacity for essential processes during this phase [62,63]. Redox-associated proteins increased in prominence but were primarily linked to NAD(P)H-dependent metabolic processes, suggesting indirect redox perturbation arising from altered metabolic flux rather than direct ROS accumulation [64,65]. This metabolic reorganization enables the pathogen to maintain energy homeostasis and biosynthetic capacity under water limitation, representing a critical transition from emergency response to sustained adaptation (Fig 4A-B, S3 Table). In contrast to the NaCl mid-phase response (6–12 HPT), which featured a transition from glutathione-based to thioredoxin-based antioxidant systems alongside metabolic restructuring, the PEG mid-phase response shows redox disturbance primarily as an indirect consequence of metabolic flux changes rather than active antioxidant system switching [17]. This mechanistic difference reflects the distinct primary challenges, salt stress demands active ionic homeostasis and ROS neutralization, while osmotic stress requires metabolic economy and selective biosynthetic prioritization. The maintenance of translational capacity in both stresses suggests a shared adaptive principle, continued selective protein synthesis is essential under both conditions, but the metabolic context differs fundamentally.

Late-stage responses (12-24 HPT) were characterized by enrichment of redox-buffering enzymes, protein quality-control systems, and membrane– or structure-associated proteins (S3 Table). The emergence of antioxidant and proteostatic functions at this stage indicates progressive accumulation of redox and protein-folding stress during prolonged exposure [66,67]. This engagement of maintenance pathways within 24h argues against acute ROS toxicity and instead supports a model in which redox buffering stabilizes metabolism and proteome integrity under sustained osmotic challenge. The temporal separation of metabolic adaptation from antioxidant responses suggests that ROS accumulation is a secondary consequence of prolonged metabolic stress rather than a primary osmotic effect, enabling *P. cinnamomi* to establish robust protective mechanisms that ensure long-term survival and rapid recovery upon rehydration. Comparative mechanistic insights shows that this delayed secondary ROS response under osmotic stress contrasts sharply with NaCl stress, where redox perturbation is primary and immediate. Under 100 mM NaCl, glutathione reductase emerged as a central regulatory hub coordinating all three phases of adaptation, whereas under PEG stress, antioxidant systems engage late and reactively as metabolic demands accumulate. This distinction illuminates why *P. cinnamomi* exhibits growth enhancement under salt stress (active metabolic mobilization in response to ionic challenge) yet maintains paradoxical growth despite widespread protein downregulation under osmotic stress (energy conservation strategy). The pathogen appears to have evolved distinct adaptive strategies, salt stress triggers offensive metabolic/antioxidant activation, whereas osmotic stress triggers defensive metabolic minimization. Both strategies converge on three-phase temporal organization, but the drivers and mechanisms are fundamentally different (Fig 4A-B, S3 Table; [17].

### Protein interaction networks reveal coordinated redox-centered regulation under PEG stress

To determine whether PEG-responsive proteins form a coordinated regulatory system, a protein-protein interaction network was constructed using STRING following homology-based mapping of *P. cinnamomi* proteins to their closest annotated homologs in *P. infestans*. Proteins passing the multiple-sample ANOVA significance threshold (FDR ≤ 0.05) were queried against the *P. infestans* proteome (STRING v12.0, strain T30-4), and interactions with a combined confidence score ≥ 0.7 were retained. The resulting network comprised 96 proteins, organized into eight temporally defined clusters (A–H) corresponding to the hierarchical clustering groups identified in Fig 4A, together with five additional first-degree interaction partners that were not assigned to a temporal cluster (Fig 4C).

Network topology revealed extensive functional connectivity among PEG-responsive proteins, indicating that the osmotic stress response is organized into integrated interaction modules rather than independent, phase-specific responses (Fig 4D, S4 Table). To determine whether individual clusters carried functional identities beyond the network-wide redox signature, STRING functional enrichment was performed separately for each of the eight temporal clusters. Clusters A and B (n = 3 proteins each) were too small to yield statistically significant enrichment and are described only by their constituent protein identities (S4 Table). For the remaining six clusters, oxidoreductase activity (GO:0016491) was the dominant enriched term in every case (FDR range 4.7 × 10 to 8.76 × 10 ¹¹), confirming redox function as a shared baseline across the network. Beyond this common signature, however, each cluster showed additional, non-overlapping enrichment reflecting distinct biochemical roles. Cluster C was specifically enriched for NADP binding (FDR = 4.0 × 10; PITG_00146, PITG_02182, PITG_02854), indicating a preference for NADPH-dependent reducing reactions, whereas Cluster F was instead enriched for NAD binding (FDR = 2.0 × 10), FMN binding (FDR = 0.0058), and phosphoglycerate dehydrogenase activity (FDR = 0.0113; PITG_01407, PITG_13165), linking this cluster to serine biosynthesis alongside redox buffering and marking a clear cofactor-preference contrast with Cluster C.

Cluster D was distinguished by enrichment for L-malate dehydrogenase activity (FDR = 0.0013; PITG_15476, PITG_21153), pointing to a role centered on TCA-cycle-linked redox balance rather than general oxidoreductase function, while Cluster H carried a markedly different signature, enriched for CoA-ligase and acid-thiol ligase activity (FDR = 0.0246 and 0.0378; PITG_00719, PITG_00720) in addition to its redox annotation, a functional class not observed in any other cluster, implicating this group in acyl-CoA and fatty-acid metabolism rather than redox chemistry alone. By contrast, Clusters E and G showed oxidoreductase activity without a comparable secondary specialization, the former alongside a general small-molecule-binding signature (FDR = 0.0194) and the latter with no additional distinguishing term. Taken together, these results indicate that although oxidoreductase function is broadly distributed across the network, the temporal clusters are differentiated by distinct cofactor preferences NADP-specific in Cluster C versus NAD-specific in Cluster F and by additional, non-redox biochemical roles, including malate dehydrogenase activity in Cluster D and CoA-ligase activity in Cluster H, indicating that these clusters represent biochemically distinct modules rather than functionally interchangeable subdivisions of a single redox response.

## Conclusion

While conducted under control *in vitro* conditions using PEG as a model osmoticum, this work reveals the proteomic mechanisms underlying *P. cinnamomi’s* ability to sustain growth under mild water limitation, an adaptive capacity that likely contributes to its persistence in dynamic environments, independently of host interactions.

Osmotic water limitation represents a recurrent challenge for soilborne pathogens in environments characterized by drought-rain cycles. By isolating osmotic stress from ionic effects, this study demonstrates that *P. cinnamomi* responds to reduced water availability through a structured, energy-saving adaptive program that preserves growth capacity rather than triggering emergency responses. The temporally organized proteomic changes reveal sequential engagement of regulatory, metabolic, and redox systems, fundamentally differing from ionic stress responses where antioxidant systems activate rapidly to combat immediate toxicity. In contrast, comparative analysis with salt stress responses (NaCl treatment) reveals that osmotic stress elicits a slower, more metabolically strategic adaptation, supporting the ecological relevance of distinguishing osmotic tolerance from ionic stress tolerance in soilborne pathogens [17].

Network analysis shows that redox-associated function is broadly distributed across every temporal cluster of the interaction network, rather than confined to a single phase or module. At the same time, cluster-level enrichment reveals that individual clusters are further distinguished by additional, non-overlapping biochemical roles beyond this shared redox baseline, including NADP-versus NAD-dependent cofactor preferences and non-redox functions such as malate dehydrogenase and CoA-ligase activity. This establishes redox regulation as a pervasive, network-wide coordinating framework for system-level adaptation under water-limited conditions, within which biochemically distinct modules perform complementary roles rather than acting as damage-response elements alone. This adaptive capacity may confer significant ecological advantages in fluctuating soil moisture conditions and informs predictions of pathogen behavior under changing climate conditions. These insights may guide targeted management strategies, including disruption of redox-buffering pathways or selective inhibition of the temporal metabolic remodeling program, to render *P. cinnamomi* vulnerable under drought conditions in water-stressed agricultural systems where osmotic stress predominates.

## ACKNOWLEDGEMENTS

We thank Rebecca McDougal and Daryl Herron from the New Zealand Bioeconomy Science Institute (Scion) for kindly providing *P. cinnamomi* samples, and Steven Whisson from the James Hutton Institute, Dundee, for valuable feedback.

## Supporting information

**S1 File: Total number of proteins identified and proteins in Venn diagrams**

**S2 File: List of proteins identified in high and low abundance**

**S3 File: Proteins identified in various clusters derived from Hierarchical clustering**

**S4 File: Network analysis of *P. infestan*s string ID homologues to identified *P. cinnamomi* proteins identified.**

